# Close homologue of L1 sensitizes lung cancer cells to cisplatin and paclitaxel via inhibition Akt pathway

**DOI:** 10.1101/747238

**Authors:** Xiangdao Cai, Bang Hu, Sheng Liu, Maolin Liu, Yunhe Huang, Peng Lei, Zhi Zhang, Zhiwei He, Linquan Zhang, Rimao Huang

## Abstract

Drug resistance is a serious promble during chemotherapy in lung cancer, which may lead to tumor relapse and further progression. CHL1 was a tumor suppressor in most malignancies, and it was found downregulated in NSCLC cisplatin-resistant cells H460. Thus, in this study, we investigated the role and mechanism of chemoresistance by CHL1 in lung cancer. Human lung adenocarcinoma cell lines A549 and its cisplatin resistant cells (A549/DDP) and paclitaxel resistant cells (A549/PTX) were applied in this research. CHL1 was found obvious downregulation in A549/DDP and A549/PTX cells versus A549 cells. Suppression of CHL1 in A549 cells, promoted cell survival rate and clone formation, decreased cell apoptosis when treated with or without DDP and PTX, respectively. While excessive CHL1 expression in A549/DDP and A549/PTX cells, the results were opposite. Moreover, CHL1 knockdown mediating chemoresistance was reversed by Akt inhibitor SC66 in A549 cells. In summary, overexpression of CHL1 reversed chemoresistance to cisplatin and PTX via suppressing Akt pathway in lung cancer, it was suggested that CHL1 maybe as a potential target for overcome chemoresistance in lung cancer.

## Introduction

Lung cancer is the most common human malignancy, accounting for 21.7% of all cancer-related deaths worldwide, and the morbidity and mortality rank the first among all malignant tumors(Chen 2015). According to the differentiation degree and morphological characteristics of cancer cells, it can be roughlt classified non-small-cell lung cancer (NSCLC) and small-cell lung cancer(SCLC). Among the lung caner, nearly 80% patients were are diagnosed as NSCLC, which manifests earlier diffusion and metastasis(Cai et al. 2018). At present, resection, chemotherapy, radiotherapy and targeted therapy are the main treatments in lung cancer. For patients with advanced NSCLC or who are physically incapacitated for surgery, chemotherapy is the preferred treatment. Cisplatin (DDP) and paclitaxel (PTX) are the first-line chemotherapeutic agents which are widely used in treating NSCLC(Han et al. 2016). Although the overall survival of patients reveived DDP or PTX based chemotherapy have been improved significantly(Liu et al. 2013), intrinsic or acquired resistance lead to tumor relapse and further progression, which have seriously limited the chemotherapy effect. Therefore, it’s crucial to explore the key molecules related to drug resistance and potential mechanisms, and that may bring a better result in lung cancer treatment.

Close homologue of L1 (CHL1) is a member of the L1 family of nerve cell adhesion molecules, and located on 3q26 locus. As a nerve cell adhesion molecule, CHL1 play an important role in the development, regeneration and plasticity of the nervous system(Long et al. 2012). The absence or mutation of CHL1 can trigger 3p syndrome and schizophrenia(Sun et al. 2012, Tassano et al. 2014). The abnormal expression of CHL1 may lead to reduced working memory and social behavior, mental damage and abnormal behavior(Morellini et al. 2007). CHL1 is mainly expressed in the central nervous system and also can be seen in other normal tissues. Recently, it was reported that CHL1 was involved in carcinogenesis and progression in a variety of human cancers. In esophageal squamous cell carcinoma (ESCC), CHL1 down-regulation was associated with invasion, lymph-node metastasis, and poor overall survival(Tang et al. 2019). The expression of CHL1 was downregulated by hypermethylation in human breast cancer, and it negative expression was contribute to breast tumorigenesis and progression(He et al. 2013, Martin-Sanchez et al. 2017). In thyroid cancer(Zhu et al. 2014), and colonic adenocarcinoma(Yu et al. 2018), CHL1 were acted as a tumor suppressor. By analyzing the Gene Expression Omnibuso (GSE21656) of NSCLC cisplatin-resistant cells H460 and parental cells, we found that CHL1 expression in cisplatin-resistant cells was significantly lower than that of parental cells, suggesting that CHL1 may be involved in NSCLC chemotherapy resistance.

In this study, we assessed the expression of CHL1 in DDP and PTX-resistant A549 cell lines and the parental cell lines, and functional studies of CHL1 were investigated to explore its potential role in regulation of chemoresistance. It was revealed that CHL1 enhance NSCLC chemosensitivity through inhibition Akt pathway. These datas showed that CHL1 may be a promising target to improve the efficacy of chemosensitivity in NSCLC.

## Results

### CHL1 is downregulated in A549/DDP and A549/PTX resistant cells

In order to explore the mechanisms of chemoresistance in lung cancer, the representative lung adenocarcinoma cell lines A549 and its cisplatin resistant cells (A549/DDP) and paclitant resistant cells (A549/PTX) were applied in this research. First, MTT assays was used to detected the cell survival rate of resistant cells when exposed to different dose of DDP and PTX, with the increased concentration of DDP (0–8 μg/ml) and PTX (0-160 ng/ml), all the cells showed a decreased survival rate, but A549/DDP and A549/PTX cells were respectively resistance to DDP and PTX compared with A549 cells (Figure 1A). The half maximal inhibitory concentration (IC_50_) values of A549 and A549/DDP cells to DDP were (1.68±0.18) and (8.30±0.92) μg/ml respectively. The IC_50_ values of A549 and A549/PTX cells to PTX were (36.97±2.56) and (174.80±8.64) ng/ml respectively (Figure 1B). Furthermore, A549/DDP and A549/PTX cells had a high expression of drug-resistant markers, MDR1 MRP and LRP compared with that of A549 cells (Figure 1C). As detected by qRT-PCR and Western blot methods, the expression of CHL1 mRNA and protein were lower in A549/DDP and A549/PTX than in A549 cells (Figure 1D and 1E). This result was consistent with the microarray analysis of H460 cispatin resistant cells (GSE21656) (Figure 1F). These results suggested that CHL1 may be involved in regulation DDP and PTX resistance in NSCLC.

**Figure 1:**
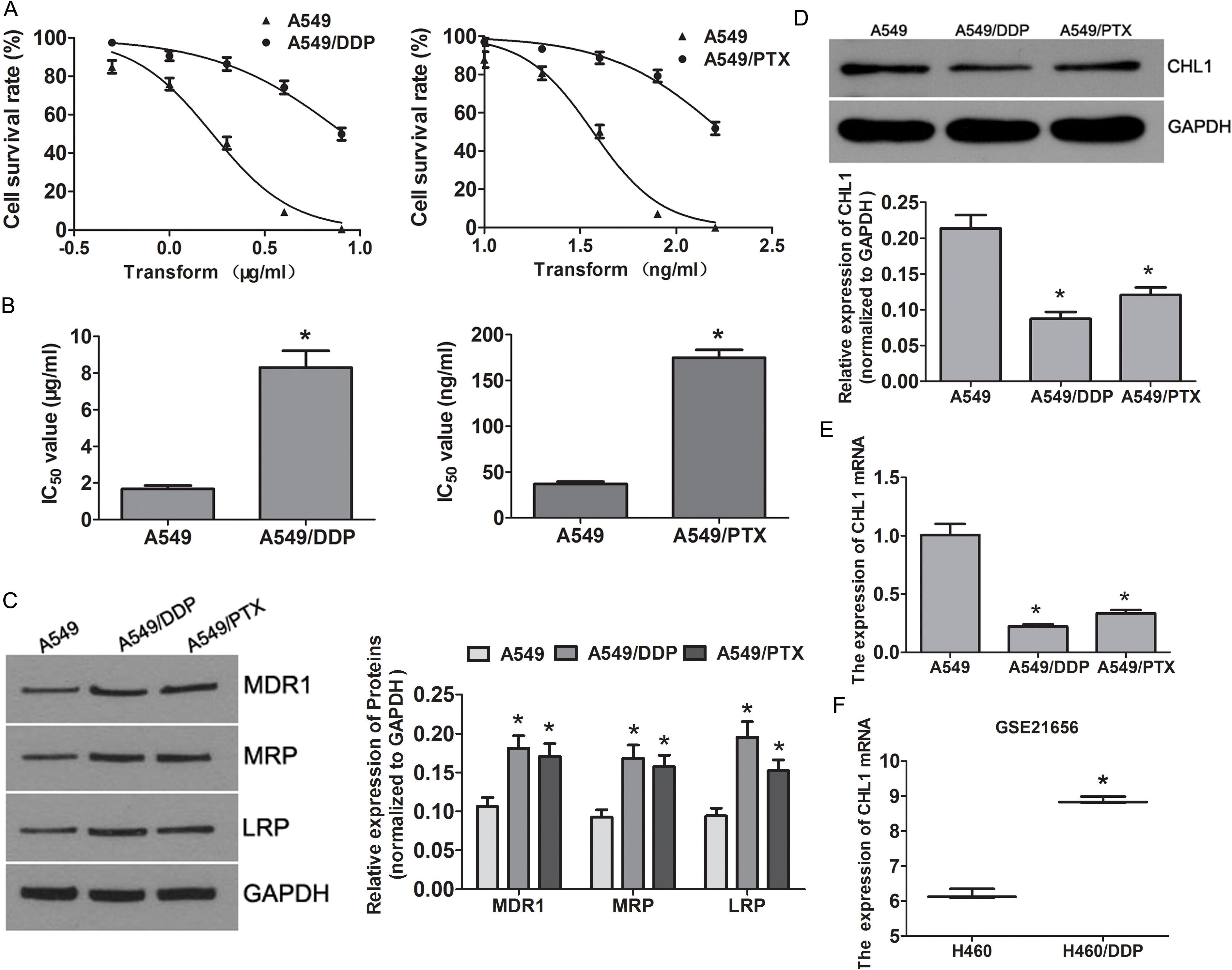
CHL1 was downregulated in DDP and PTX resistant A549 cells. (A) Cell survival of A549 and A549 resistant cells (A549/DDP, A549/PTX) treated with increasing concentration of DDP and PTX were measured by MTT assay. (B) The IC_50_ values of DDP in A549/DDP, A549 cells, and the IC_50_ values of PTX in A549/PTX, A549 cells. (C) Western blot assays dectected the expression of drug resistance-related genes MDR1, MRP and LRP in A549 cells and A549 resistant cells (A549/DDP, A549/PTX). The expression of CHL1 expression in A549 cells and A549 resistant cells (A549/DDP, A549/PTX) were analysed by Western blot (D) and qRT-PCR assays (E) respectively. (F) The expression of CHL1 mRNA in GSE21656 data. All results were expressed as mean ± SD, and were performed in triplicate. \**P* < 0.05.

### Knockdown of CHL1 resist A549 cells to DDP and PTX

There was relatively high expression of CHL1 in A549 cells as shown in above results. thus we silenced CHL1 by transfecting A549 cells with siRNAs. In the CHL1 siRNAs transfected groups, CHL1 expression was obviously reduced compared with that of the scrambled control group (Figure 2A). As siRNA-1 had a high interference efficiency, so we chose it in the follow-up experiments. When CHL1 was knockdown, it could enhanced the resistance to DDP and PTX in A549 cells (Figure 2B and 2C). Colony formation assays revealed that CHL1 silenced increased the ratio of colony formation in the absence of chemotherapeutics. Silencing of CHL1 resisted A549 cells to DDP(1.5 μg/ml) and PTX (35 ng/ml), as reflectd by a significant promoted colony formation (Figure 2D). We also explored whether CHL1 affected drug sensitivity by mediating chemotherapeutics-induced cell apoptosis. The results revealed an reduced cell apoptosis in CHL1 knockdown cells after DDP(1.5 μg/ml) and PTX (35 ng/ml) treatment (Figure 2E).

**Figure 2:**
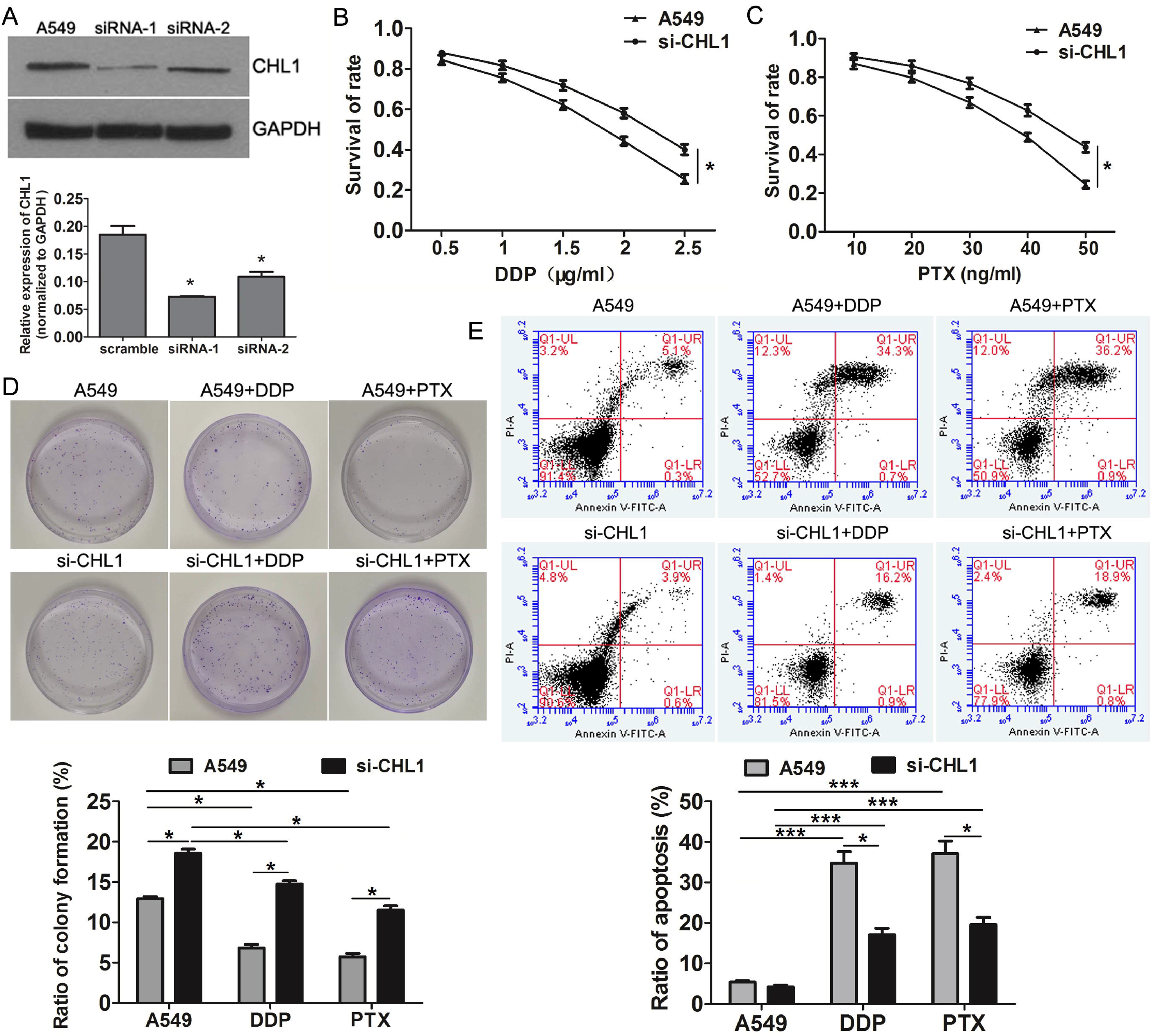
CHL1 knockdown resisted A549 cells to DDP and PTX. (A) Western blot assay was performed to detect CHL1 interference efficiency in A549 cells transfected with CHL1 siRNAs. MTT assays were performed to determine survival rate of CHL1 knockdown A549 cells treated with 0–2.5 μg/ml DDP (B) or 0–50 ng/ml DDP (C). (D) Colony formation assay of A549 cells transfected with CHL1 siRNA in the presence or absence of 1.5 μg/ml DDP and 35 ng/ml PTX. (E) Flow cytometry analysis was used to detect apoptosis in A549 cells transfected with CHL1 siRNA in the presence or absence of 1.5 μg/ml DDP and 35 ng/ml PTX. All results were expressed as mean ± SD of three replications. \**P* < 0.05, \*\*\**P* < 0.001.

### CHL1 overexpression enhances the sensitivity of A549 resistant cells to DDP and PTX

As CHL1was low expression in A549/DDP and A549/PTX cells, so we forced CHL1expression by transfected A549/DDP and A549/PTX cells with CHL1 recombinant expression plasmids (Figure 3A). MTT detection demonstrateed that the forced expression of CHL1 increased sensitivity to DDP and PTX compared with the control group (Figure 3B and 3C). Moreover, CHL1 overexpression inhibited colony formation in the absence of DDP and PTX, and led to sensitivity of resistant cells to DDP (8 μg/ml) and PTX (160 ng/ml) respectively, as shown by a significant reduced colony formation (Figure 3D). Besides, Flow cytometry results demonstrated that forced CHL1expression promoted cell apoptosis in resistant cells treated by DDP (8 μg/ml) and PTX (160 ng/ml), respectively (Figure 3E).

**Figure 3:**
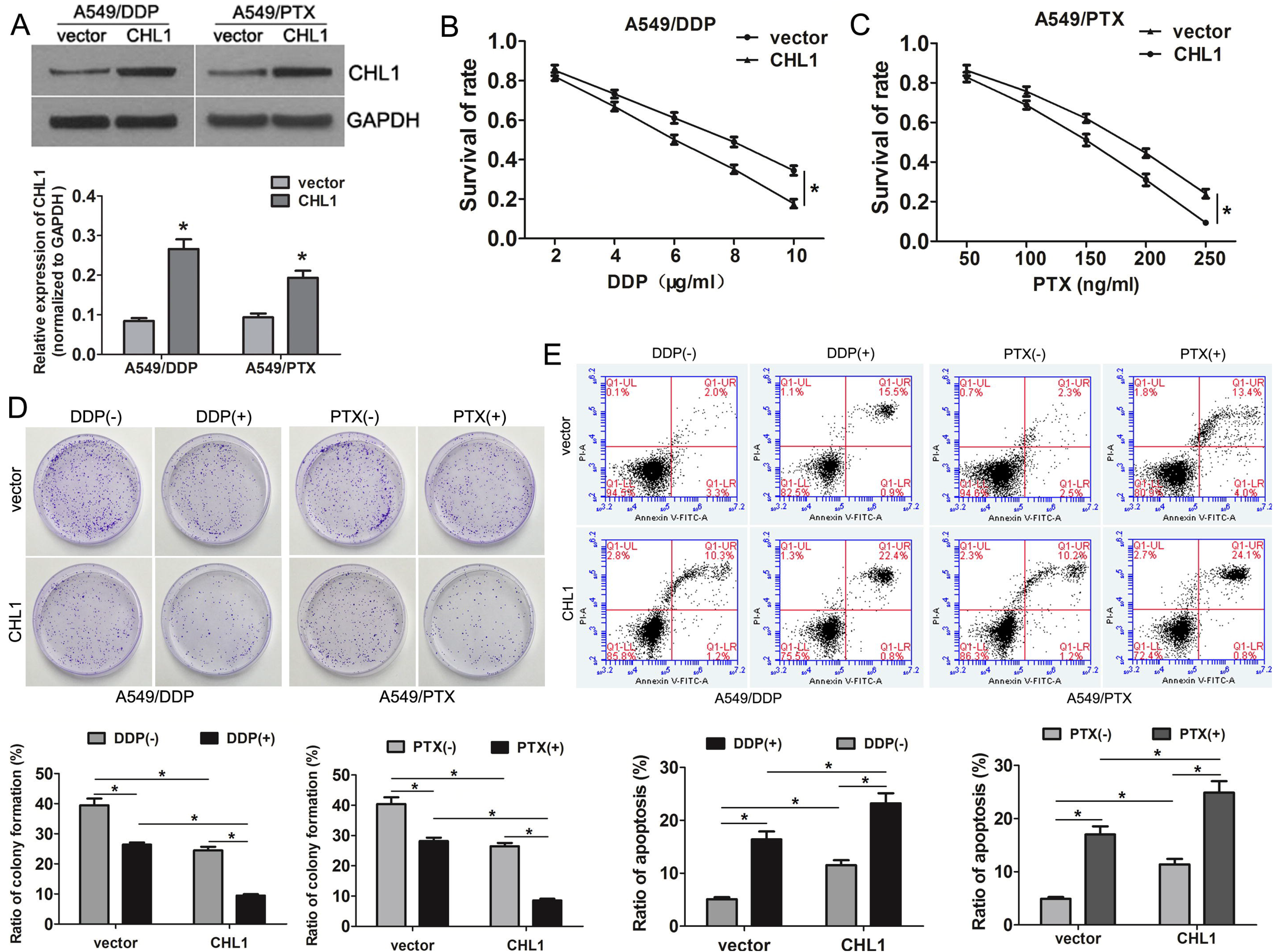
Overexpression of CHL1 enhanced the sensitivity of A549 resistant cells to DDP and PTX. (A) Western blot assay was performed to detect CHL1 expression in A549/DDP and A549/PTX cells transfected with CHL1 expression plasmids. Effect of CHL1 overexpression on A549 resistant cell survival rate when treated with 0-10 μg/ml DDP (B) or 0-250 ng/ml PTX (C). were determined by MTT assay. (D) Colony formation assays showed the number of colonies of A549 resistant cells transfected with CHL1 expression plasmids in the presence or absence of 8 μg/ml DDP and 160 ng/ml PTX. (E) Flow cytometry analysis was used to assess cell apoptosis in A549 resistant cells transfected with CHL1 expression plasmids in the presence or absence of 8 μg/ml DDP and 160 ng/ml PTX. All data represent the mean ± SD of three replications. \**P* < 0.05.

### CHL1 mediates chemosensitivity via inhibition Akt activity

Recently, research have been showed that CHL1 could inhibits Akt activity (Ognibene et al. 2018, Tang et al. 2019). To evaluate whether CHL1 mediated the chemoresistance via Akt pathway in NSCLC, we detected the expression of Akt in CHL1 silenced and overexpressed cells. In A549 cells, CHL1 knockdown promoted the expression of p-Akt^ser473^, while in A549/DDP and A549/PTX cells, restoring CHL1 expression inhibited p-Akt activation (Figure 4A), suggesting CHL1 regulation chemosensitivity may acting on Akt pathway. Furtherly, we treated the CHL1 silenced A549 cells with Akt inhibitor SC66, It was demonstrated that inhibition Akt activity might antagonized the promoted effect on cell survival (Figure 4B) and clone formation (Figure 4C), and the inhibitory effects on cell apoptosis (Figure 4D) induced by CHL1 downregulation. These results confirmed that CHL1mediates chemosensitivity in NSCLC through inhibition Akt pathway.

**Figure 4:**
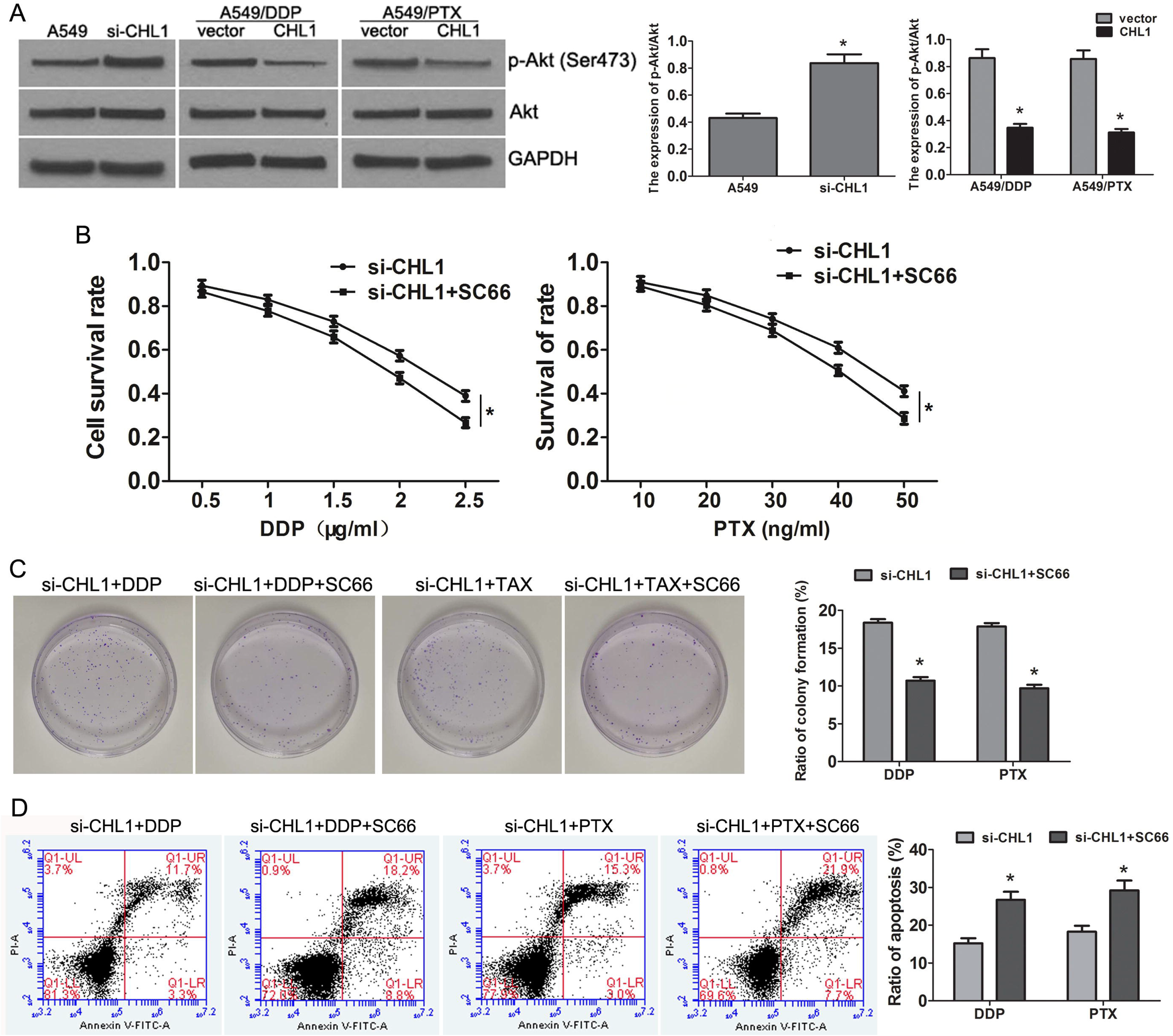
CHL1 mediates DDP and PTX sensitivity via inhition Akt activity. (A) Western blot assay was performed to detect the expression of p-Akt and total Akt in CHL1 silenced and restored cell models. (B) MTT assays were performed to detect cell survival rates of A549 cells treated with CHL1 siRNA and Akt inhibitor SC66. (D) Colony formation assays were excuted in A549 cells treated with CHL1 siRNA and Akt inhibitor SC66 in presence DDP (1.5 μg/mL) and PTX (35 ng/ml). (E) Cell apoptosis were measured in A549 cells treated with CHL1 siRNA and Akt inhibitor SC66 in presence DDP (1.5 μg/mL) and PTX (35 ng/ml). All data represent the mean ± SD of three replications. \**P* < 0.05.

## Discussion

DDP is widely applied in the treatment several malignancies, which acts broad spectrum of anti-tumor effects by inducing DNA damage and hindering DNA damage repair(Cai et al. 2018). Paclitaxel is another commonly used chemotherapeutic agent in clinic, which targets the microtubule cytoskeleton and impede cell division (Han et al. 2016, Liu et al. 2009). Most of patients have a good initial response to chemotherapy agents, but subsequent relapse is common as largely due to the emergence of drug resistance(Hassan et al. 2016). Thus, chemoresistance is consider as one of the main factors of poor prognosis in advanced NSCLC patients(Fang et al. 2018). Hence, it’s urgently necessary to explore the target and mechanism of chemoresistance in NSCLC.

CHL1 belong to the L1 family of nerve cell adhesion molecules, which was initially cloned in mice and its expression in mouse development was analyzed by Senchenko et al.(Senchenko et al. 2011). Through cell-cell interaction and mediating cell–cell and cell-matrix interactions, CHL1 has an important effect on the development, regeneration and plasticity of the nervous system(He et al. 2013). Previous reports have been demonstrated that CHL1 also participate in the carcinogenesis. In up to eleven types of tumor tissue, CHL1 was found significant downregulation(Senchenko et al. 2011). For neuroblastoma patients, low CHL1 expression predicted the poor prognosis, increased CHL1 expression suppressed tumor progression, anchorage-independent colony formation, and the growth of human tumor xenografts(Ognibene et al. 2018). In cervical cancer, CHL1 is the target of microRNAs, miR-10a and miR-590-5p promote human cervical cancer cells invasion and migration through negative regulation of CHL1(Chu et al. 2014, Long et al. 2012). In the majority tumors, CHL1 is a potential tumor suppressor gene, whose silencing is attribute to tumor growth, invasion and metastasis(He et al. 2013, Martin-Sanchez et al. 2017, Morellini et al. 2007, Tang et al. 2019, Zhu et al. 2014). However, CHL1 was reported promoting cell proliferation, metastasis and migration in human glioma(Yang et al. 2017). As for research on CHL1 and tumor chemoresistance rarely have been reported.

In this study, we minie the differentially expressed genes in NSCLC DDP-resistant cells in GEO datasets, CHL1 was found upregulation in DDP-resistant cells compare with parental cells, suggesting that CHL1 may be involved in NSCLC chemotherapy resistance. Similar study has been reported in ovarian cancer, through comparison and analysis the differential expression genes in 26 stage IIIC/IV serous ovarian adenocarcinomas, including 12 chemosensitive tumours and 14 chemoresistant tumours, the expression of CHL1 in chemotherapy-sensitive tumor tissues was higher than that in ovarian cancer tissues from drug-resistant patients, suggesting that CHL1 may help to predict the efficacy of chemotherapy for ovarian cancer(Choi et al. 2012). The aberrant methylation of CHL1 might be associated with the recurrence of colorectal cancer (CRC) patients after chemotherapy, 5-azadC treatment restored the 5-FU sensitivity in vitro, which also suggested that CHL1 may be involved in CRC chemotherapy resistance(Baharudin et al. 2017). In our result, we demonstrated that CHL1 was downregulation in A549/DDP. Considering multiple drug resistance was a common character, we applied another resistant cells-A549/TAX in this study. The results also showed that CHL1 was downregulation in A549/TAX. Upregulation the expression of CHL1, the sensitive of resistant cells to DDP and PTX were increased significantly, while downregulation CHL1 expression in parent cells A549, the results was just the opposite. This was first time confirmed that CHL1 is involved in chemotherapy sensitivity in lung cancer.

Akt/PKB (protein kinase B) is a serine/threonine protein kinase, which is activated by phosphorylation(Wu et al. 2019). As a key molecule of PI3K/Akt signaling pathway, phosphorylated Akt (p-Akt) regulates cell survival and growth, cell motility, angiogenesis, and prevents apoptosis(Ognibene et al. 2018). Moreover, Akt activation is associated with tumor chemoresistance(Clark et al. 2002, Wu et al. 2019). In our research, we found the expression of p-Akt was increased in CHL1 knockdown A549 cells, while ite expression was reduced in CHL1 overxepression A549/DDP and A549/PTX cells. When inhibited Akt activity by Akt inhibitor, the sensitivey to DDP and PTX in CHL1 knockdown A549 cells was resotred. It was verified that CHL1 enhances the sensitivity of NSCLC to chemotherapy by inhibiting Akt pathway.

In summary, we identified that CHL1 is contributed to chemosensitivity in NSCLC, and upregulation of CHL1 enhances the sensitivity of NSCLC to chemotherapy via inhibiting Akt pathway. Our results incicated that CHL1 may be a potential target to improve the efficacy of chemosensitivity in NSCLC.

## Materials and methods

### Cell culture

The human NSCLC cell lines A549, the PTX-reistant cell line A549/PTX, and the DDP-reistant cell line A549/DDP were purchased from Procell Life Science&Technology Co., Ltd (Wuhan, China). The cells were cultured in Ham’s F-12K medium (Thermo Fisher, USA) supplemented with 10% fetal bovine surum (FBS) (Thermo Fisher, USA) in a 37°C humidified incubator with 5% CO_2_.

### qRT-PCR assay

Total RNAs were isolated using Trizol reagent (Thermo Fisher, USA), and the mixed DNA was eliminated by DNase I (New England Biolabs, USA). RNA reverse transcribed with GoScript™ Kit (Promega, USA) according to the product’s instructions. CHL1 expression was measured by qRT-PCR assay using LightCycler480 system (Roche, Germany). CHL1 primers were as follows: 5’-GGCTTGGTCTCTTGCTTTCC-3’ (forward), 5’-ATCTTCCCTCCCTTTGCACG-3’ (reverse); β-actin used as the endogenous control gene for normalizing. Each reactions were conducted in triplicate.

### Cell transfection

The CHL1 recombinant expression plasmid was purchased from Sino Biological (Beijing, China). The CHL1 siRNAs were purchased from RiboBio Co., Ltd (Guangzhou, China). Cell transfection was carried out by Lipofectamine 2000 (Thermo Fisher, USA) following the manufacturers’ instructions. Cells were collected for the subsequent examination at 48 h after transfection.

### Cell viability

Cell viability was dectectd with methyl thiazolyl tetrazolium (MTT) assay. Cells were seeded in 96-well plates (1×10^4^ cells/well) with 100 μl medium per well and incubated overnight. After treated with different concentration of DDP or PTX for 48h. 100 μl MTT (5 mg/ml) solution were added to each well and incubated for 4 h at 37°C. Then the supernatant was removed and 150 μl/well DMSO was added. Subsequently, the absorbance of was measured with Microplate Reader (BioTek, USA) at 570 nm.

### Clone formation assay

Cells (1×10^3^ cells/dish) were seed into a 35 mm dish (in triplicate) and incubated overnight. Then culture media were replaced with fresh F-12K medium containing DDP or PTX for 48h. Subsequently the cells were cultured in fresh F-12K medium supplemented with 10% FBS. Two weeks later, cells were fixed in 4% paraformaldehyde and stained with 0.01% crystal violet dye for 1h at room temperature..

### Western blot

Cells were collected and washed twice with PBS. Then the cells were lysed with RIPA buffer (ThermoFisher, USA). Proteins were isolated from cell lysis buffer and equal amounts were separated by 10% SDS-PAGE gel. Afterward, the proteins were transferredto the polyvinylidene difluoride (PVDF) membrane (Thermo Fisher, USA). After blocked by 5% BSA for 2 h at 4°C, the membrane was incubated with primary antibodies anainst CHL1 (1:500, proteintech), MDR1 (1:500, proteintech), MRP (1:500, proteintech), LRP(1:500, proteintech), phospho-Akt(1:1000, Abcam), Akt (1:2000, Abcam), overnight at 4°C. After washing three time with PBS, the membrane was incubated with HRP-conjugated secondary antibody for 2 h. Finally, the immune blots were detected with enhanced chemiluminescence reagent (Thermofisher, USA).

### Flow cytometry

Cells (5×10^5^ cells/well) were seed into 6-well plates and incubated overnight. Then the cells were exposed to DDP or PTX for 48h. Cells apoptosis was measured with the FITC Annexin V Apoptosis Kit (BD Pharmingen, USA) according to the manufacturer’s instruction. Cells were collected and washed by PBS two times prior to being suspended in 500 μl binding buffer. Subsequently, cells were incubated with 5 μl Annexin V-FITC and 5 μl Propidium Iodide in dark at room temperature for 10 min. Finally the cells apoptosis were analyzed by flow cytometry (CytoFlex, Beckman, USA).

## Statistical analysis

Data were presented as mean ± SD. All the described expriments were performed at least three times. Data were analyzed with the SPSS Statistics 20.0. Student’s t test was used for statistical analysis in two groups, *P*< 0.05 was considered statistically significant.

## Data Availability Statement

The datas used to support the findings of this study are included within the article

## References

Baharudin, R., Ab Mutalib, N. S., Othman, S. N., Sagap, I., Rose, I. M., Mohd Mokhtar, N. and Jamal, R. (2017) Identification of Predictive DNA Methylation Biomarkers for Chemotherapy Response in Colorectal Cancer. Front Pharmacol, 8, pp. 47.

Cai, Y., Dong, Z. Y. and Wang, J. Y. (2018) LncRNA NNT-AS1 is a major mediator of cisplatin chemoresistance in non-small cell lung cancer through MAPK/Slug pathway. Eur Rev Med Pharmacol Sci, 22(15), pp. 4879–4887.

Chen, W. (2015) Cancer statistics: updated cancer burden in China. Chin J Cancer Res, 27(1), pp. 1.

Choi, C. H., Choi, J. J., Park, Y. A., Lee, Y. Y., Song, S. Y., Sung, C. O., Song, T., Kim, M. K., Kim, T. J., Lee, J. W., Kim, H. J., Bae, D. S. and Kim, B. G. (2012) Identification of differentially expressed genes according to chemosensitivity in advanced ovarian serous adenocarcinomas: expression of GRIA2 predicts better survival. Br J Cancer, 107(1), pp. 91–9.

Chu, Y., Ouyang, Y., Wang, F., Zheng, A., Bai, L., Han, L., Chen, Y. and Wang, H. (2014) MicroRNA-590 promotes cervical cancer cell growth and invasion by targeting CHL1. J Cell Biochem, 115(5), pp. 847–53.

Clark, A. S., West, K., Streicher, S. and Dennis, P. A. (2002) Constitutive and inducible Akt activity promotes resistance to chemotherapy, trastuzumab, or tamoxifen in breast cancer cells. Mol Cancer Ther, 1(9), pp. 707–17.

Fang, Z., Chen, W., Yuan, Z., Liu, X. and Jiang, H. (2018) LncRNA-MALAT1 contributes to the cisplatin-resistance of lung cancer by upregulating MRP1 and MDR1 via STAT3 activation. Biomed Pharmacother, 101, pp. 536–542.

Han, M. L., Zhao, Y. F., Tan, C. H., Xiong, Y. J., Wang, W. J., Wu, F., Fei, Y., Wang, L. and Liang, Z. Q. (2016) Cathepsin L upregulation-induced EMT phenotype is associated with the acquisition of cisplatin or paclitaxel resistance in A549 cells. Acta Pharmacol Sin, 37(12), pp. 1606–1622.

Hassan, W. A., Yoshida, R., Kudoh, S., Kameyama, H., Hasegawa, K., Niimori-Kita, K. and Ito, T. (2016) Notch1 controls cell chemoresistance in small cell lung carcinoma cells. Thorac Cancer, 7(1), pp. 123–8.

He, L. H., Ma, Q., Shi, Y. H., Ge, J., Zhao, H. M., Li, S. F. and Tong, Z. S. (2013) CHL1 is involved in human breast tumorigenesis and progression. Biochem Biophys Res Commun, 438(2), pp. 433–8.

Liu, J., Meisner, D., Kwong, E., Wu, X. Y. and Johnston, M. R. (2009) Translymphatic chemotherapy by intrapleural placement of gelatin sponge containing biodegradable Paclitaxel colloids controls lymphatic metastasis in lung cancer. Cancer Res, 69(3), pp. 1174–81.

Liu, Y. P., Yang, C. J., Huang, M. S., Yeh, C. T., Wu, A. T., Lee, Y. C., Lai, T. C., Lee, C. H., Hsiao, Y. W., Lu, J., Shen, C. N., Lu, P. J. and Hsiao, M. (2013) Cisplatin selects for multidrug-resistant CD133+ cells in lung adenocarcinoma by activating Notch signaling. Cancer Res, 73(1), pp. 406–16.

Long, M. J., Wu, F. X., Li, P., Liu, M., Li, X. and Tang, H. (2012) MicroRNA-10a targets CHL1 and promotes cell growth, migration and invasion in human cervical cancer cells. Cancer Lett, 324(2), pp. 186–96.

Martin-Sanchez, E., Mendaza, S., Ulazia-Garmendia, A., Monreal-Santesteban, I., Blanco-Luquin, I., Cordoba, A., Vicente-Garcia, F., Perez-Janices, N., Escors, D., Megias, D., Lopez-Serra, P., Esteller, M., Illarramendi, J. J. and Guerrero-Setas, D. (2017) CHL1 hypermethylation as a potential biomarker of poor prognosis in breast cancer. Oncotarget, 8(9), pp. 15789–15801.

Morellini, F., Lepsveridze, E., Kahler, B., Dityatev, A. and Schachner, M. (2007) Reduced reactivity to novelty, impaired social behavior, and enhanced basal synaptic excitatory activity in perforant path projections to the dentate gyrus in young adult mice deficient in the neural cell adhesion molecule CHL1. Mol Cell Neurosci, 34(2), pp. 121–36.

Ognibene, M., Pagnan, G., Marimpietri, D., Cangelosi, D., Cilli, M., Benedetti, M. C., Boldrini, R., Garaventa, A., Frassoni, F., Eva, A., Varesio, L., Pistoia, V. and Pezzolo, A. (2018) CHL1 gene acts as a tumor suppressor in human neuroblastoma. Oncotarget, 9(40), pp. 25903–25921.

Senchenko, V. N., Krasnov, G. S., Dmitriev, A. A., Kudryavtseva, A. V., Anedchenko, E. A., Braga, E. A., Pronina, I. V., Kondratieva, T. T., Ivanov, S. V., Zabarovsky, E. R. and Lerman, M. I. (2011) Differential expression of CHL1 gene during development of major human cancers. PLoS One, 6(3), pp. e15612.

Sun, Y., Zheng, S., Torossian, A., Speirs, C. K., Schleicher, S., Giacalone, N. J., Carbone, D. P., Zhao, Z. and Lu, B. (2012) Role of insulin-like growth factor-1 signaling pathway in cisplatin-resistant lung cancer cells. Int J Radiat Oncol Biol Phys, 82(3), pp. e563–72.

Tang, H., Jiang, L., Zhu, C., Liu, R., Wu, Y., Yan, Q., Liu, M., Jia, Y., Chen, J., Qin, Y., Lee, V. H., Luo, S., Wang, Q. and Guan, X. Y. (2019) Loss of cell adhesion molecule L1 like promotes tumor growth and metastasis in esophageal squamous cell carcinoma. Oncogene, 38(17), pp. 3119–3133.

Tassano, E., Biancheri, R., Denegri, L., Porta, S., Novara, F., Zuffardi, O., Gimelli, G. and Cuoco, C. (2014) Heterozygous deletion of CHL1 gene: detailed array-CGH and clinical characterization of a new case and review of the literature. Eur J Med Genet, 57(11-12), pp. 626–9.

Wu, Y. H., Huang, Y. F., Chen, C. C. and Chou, C. Y. (2019) Akt inhibitor SC66 promotes cell sensitivity to cisplatin in chemoresistant ovarian cancer cells through inhibition of COL11A1 expression. Cell Death Dis, 10(4), pp. 322.

Yang, Z., Xie, Q., Hu, C. L., Jiang, Q., Shen, H. F., Schachner, M. and Zhao, W. J. (2017) CHL1 Is Expressed and Functions as a Malignancy Promoter in Glioma Cells. Front Mol Neurosci, 10, pp. 324.

Yu, W., Zhu, K., Wang, Y., Yu, H. and Guo, J. (2018) Overexpression of miR-21-5p promotes proliferation and invasion of colon adenocarcinoma cells through targeting CHL1. Mol Med, 24(1), pp. 36.

Zhu, H., Fang, J., Zhang, J., Zhao, Z., Liu, L., Wang, J., Xi, Q. and Gu, M. (2014) miR-182 targets CHL1 and controls tumor growth and invasion in papillary thyroid carcinoma. Biochem Biophys Res Commun, 450(1), pp. 857–62.

